# Culture volume influences the dynamics of adaptation under long-term stationary phase

**DOI:** 10.1101/2020.08.13.249599

**Authors:** Jonathan Gross, Sarit Avrani, Sophia Katz, Ruth Hershberg

**Affiliations:** Rachel & Menachem Mendelovitch Evolutionary Processes of Mutation & Natural Selection Research Laboratory, Department of Genetics and Developmental Biology, the Ruth and Bruce Rappaport Faculty of Medicine, Technion-Israel Institute of Technology, Haifa 31096, Israel; The Department of Evolutionary and Environmental Biology and the Institute of Evolution, University of Haifa

## Abstract

*Escherichia coli* and many other bacterial species, which are incapable of sporulation, can nevertheless survive within resource exhausted media by entering a state termed long-term stationary phase (LTSP). We have previously shown that *E. coli* populations adapt genetically under LTSP in an extremely convergent manner. Here we examine how the dynamics of LTSP genetic adaptation are influenced by varying a single parameter of the experiment - culture volume. We find that culture volume affects survival under LTSP, with viable counts decreasing as volumes increase. Across all volumes, mutations accumulate with time, and the majority of mutations accumulated demonstrate signals of being adaptive. However, positive selection appears to affect mutation accumulation more strongly at higher, compared to lower volumes. Finally, we find that several similar genes are likely involved in adaptation across volumes. However, the specific mutations within these genes that contribute to adaptation can vary in a consistent manner. Combined, our results demonstrate how varying a single parameter of an evolutionary experiment can substantially influence the dynamics of observed adaptation.

## Introduction

*Escherichia coli* is capable of surviving for months and even years within batch culture, without the addition of any external growth resources, in a state termed long-term stationary phase (LTSP)^1–7^. When placed into fresh rich media, *E. coli* will undergo a short period of exponential growth, followed by a short stationary phase and a rapid death phase. Death phase does not, however, lead to the extinction of the entire *E. coli* population. Instead, populations can enter LTSP and maintain viability over many months and even years.

We have previously published a paper describing the dynamics of adaptation during the first four months spent under LTSP^7^. We found that *E. coli* LTSP populations adapt genetically through the rapid acquisition of mutations. These mutations are enriched for functional categories, relative random expectations indicating that positive selection strongly affects their accumulation. Further evidence for the adaptive nature of mutation accumulation is the extreme convergence with which mutation accumulation occurs. The most striking example of convergent mutation accumulation under LTSP involves the RNA polymerase core enzyme (RNAPC). We found that across all populations and time points sampled ∼90% of the hundreds of sequenced clones carry a mutation within one of two of the genes encoding the RNAPC, *rpoB* and *rpoC*. For >90% of the clones carrying an RNAPC mutation, we found that one of only three specific sites was mutated: RpoB position 1272, RpoC position 334 or RpoC position 428. Clones containing mutations within each of these positions appeared within all five independently evolving LTSP populations. Such high levels of convergence demonstrate that RNAPC mutations and particularly mutations to these three specific sites of the RNAPC are adaptive within our LTSP populations.

Previous studies into LTSP that did not sequence whole genomes, suggested that *E. coli* adaptation under LTSP often relies on mutations within the master regulator of the stress response, RpoS^2,5,8,9^. It was therefore surprising that we saw no mutations within RpoS in our LTSP populations. One possibility for this apparent discrepancy was that the conditions under which LTSP was studied in our experiments varied from those of previous studies. While most past studies examined adaptation to LTSP as it occurs in much smaller volumes (≤ 5ml) in test tubes, we examined adaptation to LTSP in large volumes (400 ml) within flasks. Indeed, it has been demonstrated that culture volume and receptacle, as well as media type can affect survival dynamics under LTSP^10–14^. Culture volume in particular can affect both the overall population size and the conditions faced by the bacteria, prior to and during LTSP.

In the current study, we examine how varying culture volume affects the dynamics of genetic adaptation under LTSP. We carry out LTSP experiments, followed by whole-genome sequencing of individual evolved clones within two additional volumes: 40 ml and 4 ml, while maintaining all other experimental parameters consistent with our previous 400 ml experiments. Our results suggest that manipulation of this one parameter can substantially affect the dynamics of adaptation as these pertain to rates of mutation accumulation, the influence of natural selection on mutation accumulation, and the identity of loci and specific sites involved in adaptation.

## Materials and Methods

### LTSP evolutionary experiments growth conditions

All our experiments were carried out in flasks of the same shape, with the same relative ratio between growth media and flask sizes. To initiate a new LTSP evolutionary experiment, a single colony of *E. coli* K12 MG1655 was grown overnight. Single colonies were used to inoculate test tubes with 4ml of fresh Luria Broth (LB) medium. Each culture was then grown until it reached an OD (600nm) of 0.4, and then ∼2 × 10^9^ cells (1 ml) were used to inoculate 4 ml, 40 ml and 400 ml of LB in 20 ml glass flask and 200 ml and 2,000 ml polycarbonate breathing flask, respectively. This procedure was used in order to both start with a similar number of cells in all populations and start with isogenic populations by reducing the generations passed since each population was a single cell. The ten resulting flasks were placed in an incubator set at 37 °C, shaking at 225 rpm. No new nutrients or resources were added to the cultures with time, except for sterile water that was added to compensate for evaporation every day, according to the weight lost in each flask during that time.

### Sampling LTSP Populations and Estimating Viability

One ml of each culture was sampled, initially, every day, then every week, then every month and following that, at extended intervals **(fig. 1A)**. Dilutions were plated using a robotic plater to evaluate viability using live counts. Samples were frozen in 50% glycerol in a -80°C freezer for future analyses. For each sample, pH was measured using pH indicator strips.

**Figure 1.**
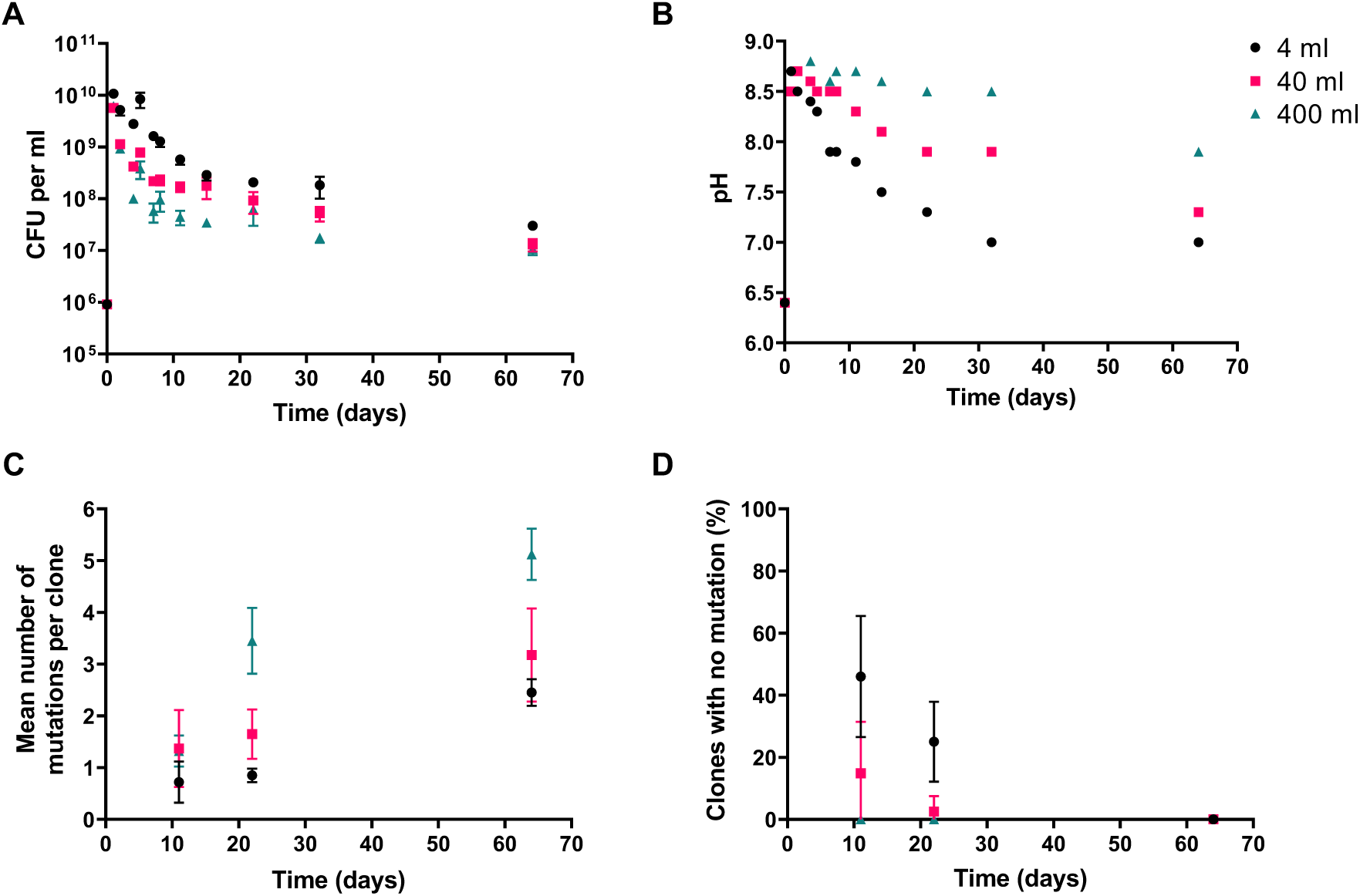
Patterns of viability, pH and mutation accumulation under LTSP vary with volume. (A) Higher viability is maintained under LTSP at lower volume populations. Depicted are the mean colony forming unit (CFU) calculations across five independently evolving populations, at each volume. Error bars represent the standard deviation around these means (B) Differences in pH are observed between volumes (C) Higher mutation accumulation under LTSP, at higher volumes. Depicted are the mean numbers of mutations accumulated at each time point. Error bars represent the standard deviation around these means (D) Higher fraction of clones survive longer, without acquiring any mutations at lower compared to higher volumes. Depicted are the mean number of clones per population that do not carry any mutations, relative to their ancestral genotype. Error bars represent the standard the deviation around these means

### Sequencing of LTSP Clones

We sequenced LTSP evolutionary experiments at two different growth volumes of 4 ml and 40 ml and analyzed them in comparison to the 400 ml growth volume, LTSP experiments previously established in our lab^7^. Frozen cultures of each population and time point were thawed and dilutions were plated and grown overnight. ∼10 clones from four of the 4 ml and 40 ml populations (eight populations in total) were sequenced at each of three time points of the experiments: 11, 22 and 64 days (**Table S1**). ∼10 clones from the fifth population of each of the two volumes were sequenced only from day 11 of the experiments (Table S1). Each colony was inoculated into 4ml medium in a test tube and were grown until they reached an OD of ∼1. Again, this was done to reduce the time these colonies can evolve and accumulate mutations. 1ml of the culture was centrifuged at 10,000 g for 5 min and the pellet was used for DNA extraction. The remainder of each culture was then archived by freezing in 50% of glycerol in a -80°C freezer. DNA was extracted using the Qiagen DNeasy Blood & Tissue Kit. Library preparation followed the protocol outlined by Baym et al.^15^. Sequencing was carried out at the Technion Genome Center using an Illumina HiSeq 2500 machine. Clones extracted from each time point were sequenced using paired-end 150 bp reads. The ancestral clones, which were used to initiate all the other populations, were sequenced as well.

### Calling of Mutations

In order to call mutations, the reads obtained for each LTSP clone or ancestral strain were aligned to the *E. coli* K12 MG1655 reference genome (accession NC_000913). LTSP clone mutations were then recorded in case they appeared within an LTSP clone’s genome, but not within the ancestral genome. Alignment and mutation calling were carried out using the Breseq platform, which allows for the identification of point mutations, short insertions and deletions, larger deletions, and the creation of new junctions^16^.

### Calculating the Proportion of Non-Synonymous Versus Synonymous Mutations Expected Under Neutrality

The DNA sequences of all protein-coding genes in *E. coli* strain MG1655 were downloaded from the NCBI database. For each position of each gene, we examined the likelihood that a mutation at this position would lead to a non-synonymous or a synonymous change. For example, any change to the third codon position of a 4-fold degenerate codon would be synonymous, so mutations at such a position would be 100% likely to be synonymous. In contrast, mutations to the third codon position of a 2-fold-degenerate codon will be synonymous for a third of possible mutations and non-synonymous for two-thirds of possible mutations. In such a manner, we could add up the likelihood of a mutation being synonymous or non-synonymous across the entirety of *E. coli* K12. MG1655’s protein-coding genes.

### Exponential Growth Rate Estimation

In order to calculate the exponential growth rate of a specific LTSP clone, single colonies were used to inoculate test tubes with 4ml of fresh LB. These cultures were grown overnight, and in the morning, they were diluted to an OD of ∼0.2. Then the cultures were allowed to grow for an additional hour and diluted again to an OD of ∼0.2 and transferred to a 96 well plate. This plate was incubated, and OD (600 nm) was read by the plate reader every 10 minutes. The exponential growth rate was calculated from the slope of the growth at the exponential phase. We used the maximum rate looking at seven consecutive time points, where R^2^>0.99. For each clone, we tested two colonies with two technical replicates.

## Results

### Culture Volume Affects survival under LTSP

In order to examine how culture volume affects the dynamics of adaptation under LTSP, we carried out LTSP evolutionary experiments, as we had previously done at 400 ml, but using culture volumes of 400 ml, 40 ml and 4 ml. For each of the volumes, five independent populations were established. All the populations were grown in flasks that can contain five times the LB volume used (two-liter flasks for 400 ml experiments, 200 ml flasks for 40 ml experiments, and 20 ml flasks for 4 ml experiments). To initiate each of the ten new populations, we inoculated ∼10^6^ cells per ml of the *E. coli* lab strain, K12 MG1655, into the relevant LB volume. From each population, across all experiments, a sample was drawn at first daily, then weekly and eventually monthly, up to day 64. No new resources were added to the cultures throughout the experiment besides distilled water to counteract culture evaporation (*Materials and Methods*). We quantified the number of viable cells at each sampling time point, by plating appropriate dilutions onto LB plates and counting the resulting colonies (**Figure 1A**). The remainder of each sample was frozen.

As shown in **Figure 1A**, across all the experiments, bacteria reached approximately the same maximal population size per ml of LB over the first 24 hours of the experiment. However, the later decline in viability during Death phase was more rapid at higher, compared to lower volumes and lower volume populations maintained higher viability upon entry into, and during LTSP. In addition to measuring CFUs at each sampling point, we also measured pH (**Figure 1B**). Although the alkaline peak (at 24 hours) of all groups was similar (∼8.6), the reduction in pH levels slowed down with volume increase. These results are consistent with previous studies showing slower death within smaller volume LTSP populations and an association between alkalization of growth media and death dynamics during death phase^8,10,17–21^.

### Lower mutation accumulation within smaller volume LTSP populations

For four of the five 40 ml populations and four of the five 4 ml populations, we sequenced ∼10 clones per population, at each of three time points: day 11, day 22 and day 64. For the fifth population of each volume, we sequenced ∼10 clones only from the day 11 time point (**Table S1**). The ancestral clones from which each population was established were also sequenced, allowing us to identify mutations that arose within each population, during its evolution under LTSP. Mutations were called using the breseq pipeline^22^. A full list of all identified mutations can be found in **Table S2**. For convenience, this table also contains the mutations identified in clones sequenced from these three time points from the five 400 ml populations, and already reported on in Avrani et al.^7^.

In three of the five 400 ml populations, we found the emergence of mutators, suffering a mutation within a mismatch repair gene^7^. These mutators accumulate a higher number of mutations relative to non-mutators. No such mutators were found in any of the ten lower volume populations. Excluding any mutators found at 400 ml, we compared the numbers of mutations accumulated at day 11, 22 and 64 between populations grown at different volumes. We found that mutation accumulation increased with volume (**Figure 1C**).

Another way to consider the accumulation of mutations is to ask what proportion of sequenced clones maintain an entirely ancestral genotype (by acquiring no mutations under LTSP). By day 64, across all volumes, we no longer see clones that carry an entirely ancestral genotype (**Figure 1D, Tables S1** and **S2**). Within the 400 ml populations, very few ancestral genotype clones are seen at day 11, and no such clones at all are seen by day 22. At the same time, cells carrying no mutations are able to survive at observable frequencies for longer periods of time within the lower volume LTSP populations (**Figure 1D**). Ancestral genotype clones are maintained at higher frequencies at days 11 and day 22 within the 4 ml, compared to the 40 ml populations.

### The majority of mutations accumulated across all volume are likely adaptive

The ratio of non-synonymous to synonymous mutations (dN/dS) is significantly higher than 1 for mutations acquired across all examined volumes (for the 400 ml populations we cannot calculate dN/dS as there are no synonymous mutations at all and so dS=0, **Table 1**). The enrichment in non-synonymous substitutions relative to random expectations is statistically significant for all volumes (*P* << 0.001, according to a χ^2^ test, Table 1 Materials and Methods). Such an enrichment in non-synonymous substitutions is considered to be a signal of positive selection affecting accumulated mutations^23^ and therefore indicates that a substantial proportion of the mutations accumulated across volumes are likely adaptive. The highest fraction of non-synonymous substitutions is found at 400 ml and the lowest at 4 ml. This might suggest that positive selection acts more strongly at higher compared to lower volumes. However, differences in the proportion of non-synonymous to synonymous substitutions are only significant when comparing the 400 ml substitutions to those accumulated at 4 ml (*P* = 0.04, according to a χ^2^ test, Table 1)

**Table 1.** Enrichment in non-synonymous substitutions relative random expectations across all culture volume sizes

| Culture volume | # non-synonymous mutations | # synonymous mutations | dN/dS<sup>a</sup> | P-value<sup>b</sup> |
| --- | --- | --- | --- | --- |
| 4 ml | 52 | 2 | 8 | <<0.001 |
| 40 ml | 146 | 2 | 22.4 | <<0.001 |
| 400 ml | 213 | 0 | N/A | <<0.001 |

<sup>a</sup>dN/dS (the ratio of the rates of non-synonymous to synonymous mutations) was calculated as
are given in the previous two columns of the table and the number of synonymous and non-synonymous sites are calculated based on a combination of all *E. coli* protein-coding genes (see Materials and Methods). dN/dS could not be calculated for the 400 ml populations as no synonymous mutations were observed. It can be said, however, to be much higher than 1.
<sup>b</sup> $\chi^2$ P-value with which it is possible to reject the null hypothesis that there is no enrichment in non-synonymous or synonymous mutations, relative random expectations based on the number of non-synonymous and synonymous sites within *E. coli* protein-coding genes.

That mutation accumulation is dominated within all volume populations by positive selection is also evident from the immense convergence with which mutations accumulate across volumes (**Figure 2**). 30, 16 and 15 loci (genes or intergenic regions) are mutated in two or more of the five populations within 400 ml, 40 ml and 4 ml, respectively. The 30 loci, which are convergently mutated across two or more of the five 400 ml populations constitute only 0.86% of the *E. coli* genome. Yet, 84.9% of the mutations found at days 11, 22 and 64 within these populations (excluding mutators) occur within these 30 loci. Similarly, the 16 and 15 loci convergently mutated within the 40 and 4 ml populations constitute only 0.54% and 0.46% of the *E. coli* genome, respectively. However, 80.5% of mutations found within the 40 ml populations and 75% of mutations found within the 4 ml populations occur within these convergently mutated loci. Such enrichment in mutations occurring within the same loci across independent populations constitutes a strong signal of positive selection. The fact that the majority of mutations found within clones evolving under LTSP occur within convergently mutated loci further indicates that the majority of mutations accumulated under LTSP are likely adaptive.

**Figure 2.**
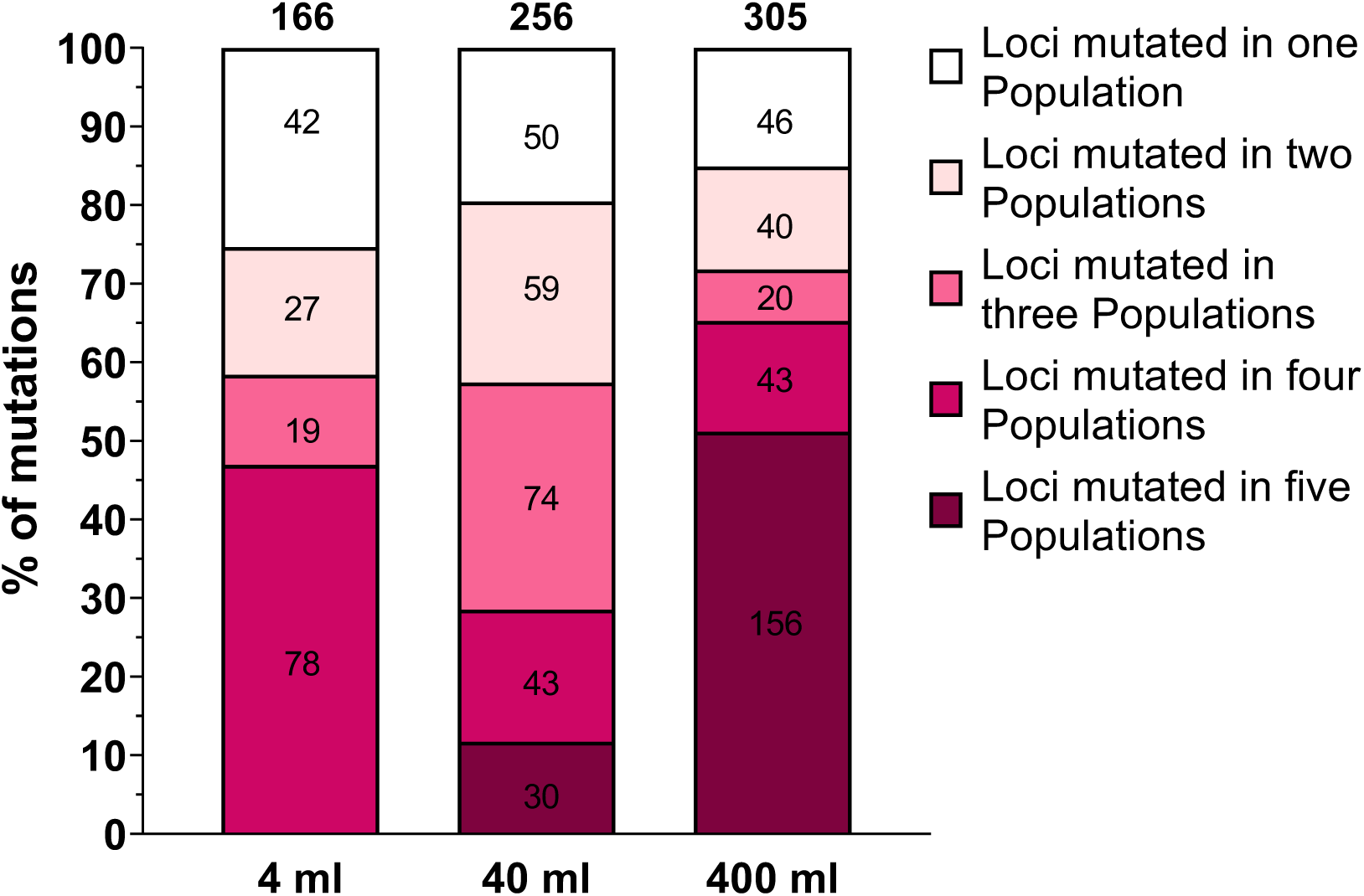
Convergent patterns of mutation accumulation observed across volumes under LTSP. Such convergence is highest at 400 ml and lowest at 4 ml. For each population, the proportion of mutations falling within genes mutated in one, two, three, four, or all five populations is presented.

While most mutations tend to fall within convergently mutated loci across all volumes, this trend appears to be more pronounced within the 400 ml populations. In other words, mutations occurring under LTSP at 400 ml tend to more often occur within loci that are mutated across a larger number of independent populations (*P* << 0.001 for comparisons against both 4 ml and 40 ml, according to a χ^2^ test, **Figure 2**). While a higher fraction of the mutations falling within the 40 ml populations falls within convergently mutated genes (80.5%) compared to mutations falling at 4 ml (75%), this difference is not statistically significant (*P* = 0.1). Combined, these results suggest that while positive selection affects a majority of LTSP mutations across volumes, a higher fraction of mutations accumulated at larger volumes are likely adaptive.

### Similar genes are often involved in adaptation under LTSP across volumes, yet specific adaptive mutations can vary according to volume

While some loci are mutated in a convergent manner in only one volume, there is substantial overlap in the identity of the loci which are convergently mutated across populations, across the different volumes (**Figure 3**). This suggests that similar genes are often involved in adaptation to LTSP across volumes. For example, across the three volumes, the genes *rpoB* and *rpoC* encoding the RNA polymerase core enzyme (RNAPC) are mutated in multiple populations. Similarly, the RNA chaperone gene *proQ*^24^, the proline metabolism genes *putA*^25^ and *putP*^26^ and the serine/threonine transporter gene *sstT*^27^ are all mutated convergently, across all three volumes. Additionally, multiple 400 ml and 40 ml populations carry mutations within the short-chain fatty acid metabolism regulator genes *fadR*^28^ and *atoS*^29^, the *paaX*^30^ transcriptional regulatory gene, the protease gene *clpA*^31^ and within the promoter region of the *cycA*^32^ transporter gene. On the whole, more convergently mutated genes are shared between the 40 ml and 400 ml populations than between either of these volume populations and the 4 ml populations. This could suggest that selective pressures might be more similar between 400 ml and 40 ml, but it could also be influenced by the fact that the 4 ml populations accumulated the lowest numbers of mutations overall (**Figure 1C**).

**Figure 3.**
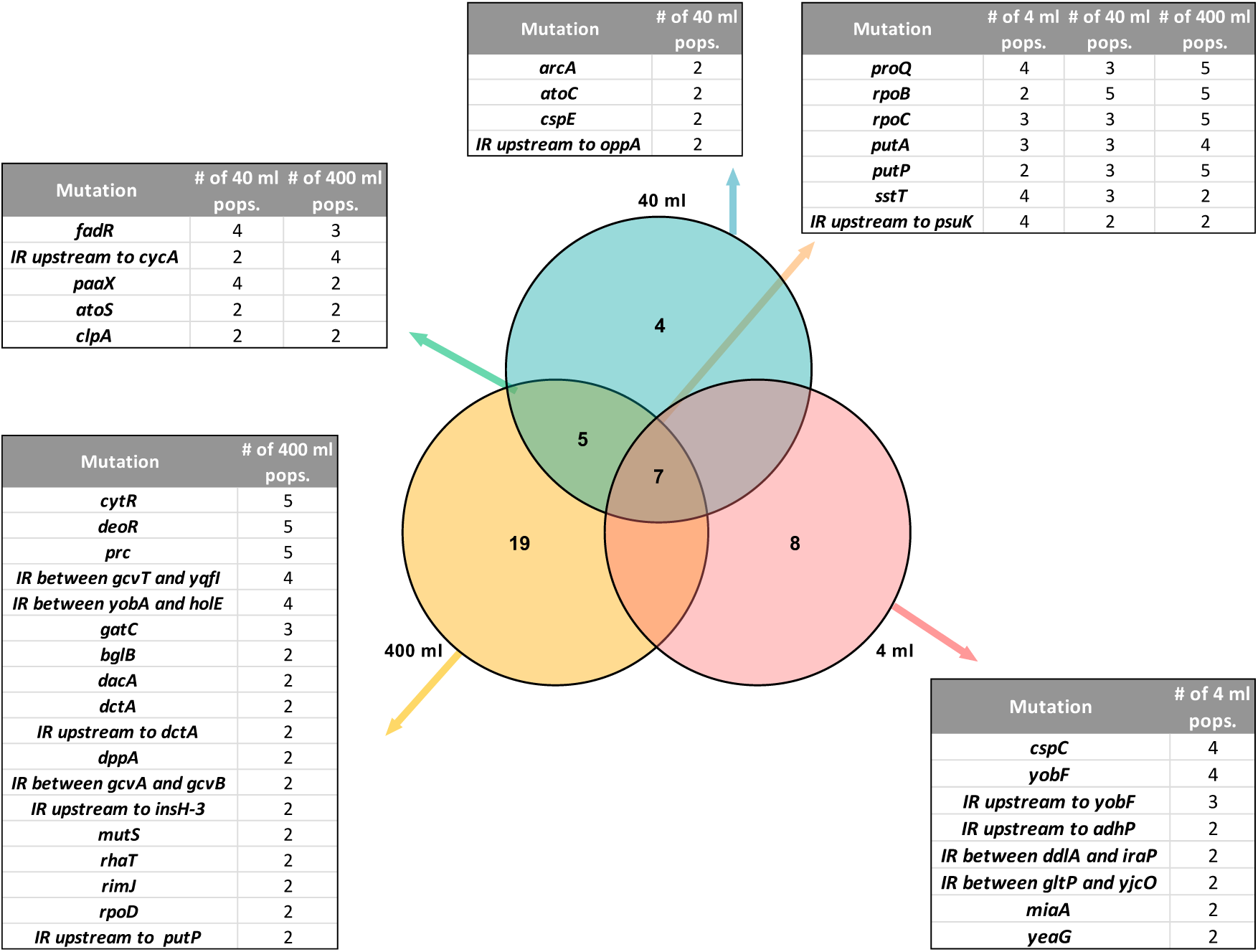
Overlap in the identity of the loci that are mutated in a convergent manner across LTSP populations, at different volumes. Presented is a Venn diagram representing the numbers of loci that are convergently mutated across multiple volumes or across only a single volume. Adjacent tables provide details on the identity of the loci presented in each section of the Venn diagram and on the numbers of populations they are mutated in. (IR=Intergenic region)

Mutations within the RNAPC genes *rpoB* and *rpoC* constituted the most striking example of convergent adaptation observed at 400 ml. In our previous study we reported that over 90% of 400 ml LTSP clones carried mutations within the RNAPC. We further reported that across all five 400 ml populations these mutations most frequently fell within only three specific sites of the RNAPC. Intriguingly, we find that at 40 ml there is also a very strong tendency for mutations to occur within specific sites of RpoB and RpoC proteins, across independently evolving populations. However, the identity of the protein sites at which mutations occur in a convergent manner is different between the 40 ml populations and the 400 ml populations (**Figure 4**). These results suggest that while mutations within the RNAPC are involved in adaptation to LTSP across volumes, culture volume may affect the identity of the specific sites of the RNAPC which are involved in this adaptation.

**Figure 4.**
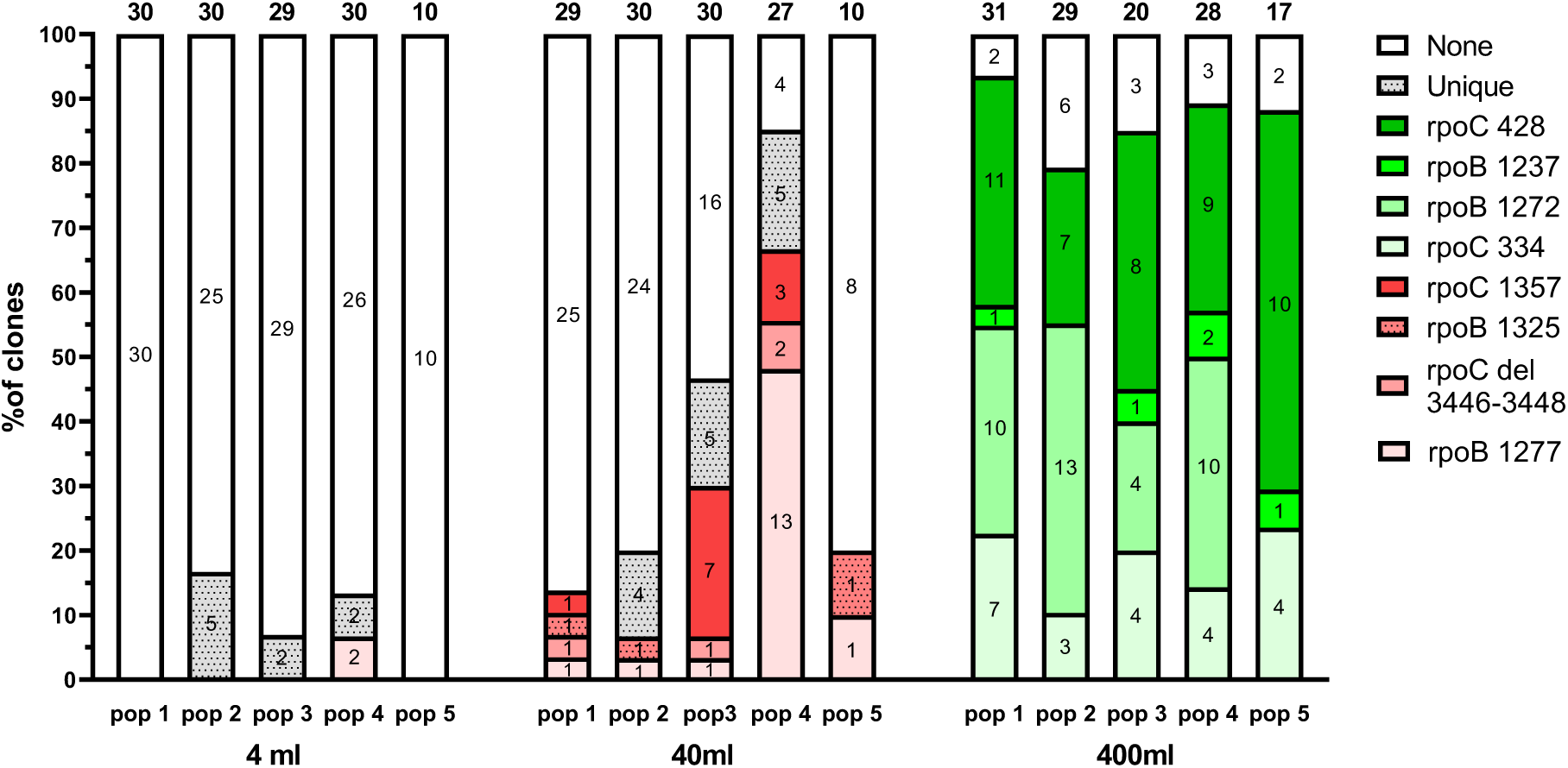
Highly convergent patterns of mutation accumulation within the RNA polymerase core enzyme. Different sites of the RNAPC are convergently mutated at different volumes. A higher fraction of clones within each population carry an RNAPC adaptation, at higher, compared to lower volumes. Depicted for each population, the proportion of clones carrying mutations within each mutated RpoB and RpoC position. Each bar represents a population and all five populations are divided by their volumes. Numbers above bars represent the total number of clones sequenced from these populations at the three time points studied. Numbers within sections of each bar represent the number of clones found to carry mutations within the site represented by that section in that population. Sites of the RNAPC that are mutated in only a single population (out of the 15 populations sampled across volumes) are binned together and designated as ‘Unique’.

While clones that do not carry an RNAPC adaptation appear to be quite rare within the 400 ml populations, they are much more frequent within the 40 ml populations. This corresponds well to our result presented in **Figure 1D**, showing that as culture volume goes down, more cells are able to survive without acquiring mutations. In other words, while the patterns of high convergence in RNAPC mutations suggest that these mutations are important for LTSP adaptation at both 40 ml and 400 ml, it appears that more cells are able to manage for longer, under LTSP, without these adaptations at the lower culture volumes (**Figure 4**).

## Discussion

Our results show that culture volume affects mutation accumulation under LTSP. Lower mutation accumulation within smaller volume LTSP populations may stem from differences in population sizes between the volumes. During the first day of growth under LTSP, across all volumes, cell numbers increase to ∼10^10^ cells per ml of culture. This means that the overall number of cells present within the 400 ml populations is about ten times higher than for the 40 ml populations and about 100 times higher than for the 4 ml populations. This might affect the number of mutations that occur within each population by that time point, which could affect the extent of standing genetic variation available for the initial adaptation. However, the fact that mutations accumulate with time under LTSP suggests that not all genetic adaptation arises from this initial standing variation. Instead, it appears that mutations continue to occur under LTSP. At later time points, the differences in population sizes between the different volumes reduce due to the fact that at lower volumes, larger numbers of cells per ml are present within the populations (**Figure 1A**).

A second possible, non-mutually exclusive, explanation for differences in mutation accumulation between the various volume populations relates to the intensity of natural selection in favor of acquiring adaptations. It is quite likely that when a smaller fraction of cells survive to enter and persist under LTSP, competition between cells for survival may be stronger. As a result, it is possible that selection to carry an adaptation enabling better survival and/or growth under LTSP may be stronger under conditions imposing a stronger constraint on the number of surviving cells. Fitting with this, we observed that within smaller volume populations, that maintain higher viability **(Figure 1A**), a higher fraction of clones persists longer, without the acquisition of any mutations (**Figure 1D**).

Patterns of mutation accumulation at different volumes also suggest that positive selection may be stronger at larger volumes. While we show that across all volumes, there is a significant enrichment in non-synonymous vs. synonymous substitutions, relative to neutral expectations, this enrichment seems to be strongest at 400 ml and weakest at 4 ml (**Table 1**). Additionally, while across all volumes a majority of mutations tend to fall within convergently mutated loci, this trend is again strongest at 400 ml and weakest at 4 ml (**Figure 2**).

Mutators that acquired a mutation within a mismatch repair gene, resulting in increased mutation accumulation, emerged in three of the five 400 ml LTSP populations by day 64. The absence of such mutators within any of the 4 ml or 40 ml populations, may also be the result of weaker positive selection affecting mutation accumulation within these populations. Mutators often emerge during evolutionary experiments and were also observed within natural bacterial populations and clinical antibiotic resistant isolates^33–38^. It is thought that mutators increase in frequencies due to indirect selection in favor of linked adaptations that occur more rapidly when overall mutation rates are higher^39–41^. Because mutators also suffer a cost due to the more rapid occurrence of deleterious mutations, mutators are thought to increase in frequency more under conditions in which there is a strong pressure to adapt^39–41^. The fact that we observe mutators only at 400 ml and not at 40 ml or 4 ml may, therefore, indicate once more that the need to adapt is stronger at higher volumes.

There is substantial overlap between volumes in the identity of the loci, which are mutated in a convergent manner. At the same time, there are also examples of loci that are mutated in a convergent manner at one volume, but not within others. More overlap is seen between the convergently mutated loci found at 400 and 40 ml, compared to the overlap observed between either 400 ml and 40 ml and the 4 ml populations (**Figure 3**). This may partially stem from the occurrence of fewer mutations overall within the 4 ml populations. However, such lower mutation accumulation within the 4 ml populations cannot explain this entire pattern. For example, mutations within the genes *cspC* and *yobF* are found in four of the five 4 ml populations but do not occur in a convergent manner at 40 ml or 400 ml. Our results, therefore, suggest that while many of the same loci may be involved in adaptation to LTSP across volumes, other loci may be involved in adaptation only within specific volume LTSP populations.

The most striking example of convergent adaptation we observe involves mutations falling within the *rpoB* and *rpoC* genes that encode the RNA polymerase core enzyme (RNAPC). Mutations within *rpoB* and *rpoC* are observed in all 400 ml and 40 ml LTSP populations, and in three of the five 4 ml LTSP populations. The majority of the RNAPC mutations occurring within the 400 ml populations fall within one of only four specific sites of the enzyme complex (**Figure 4**). Three of these sites are mutated across all five 400 ml LTSP populations. The remaining site is mutated in four of the five populations. Such extreme convergence strongly suggests that mutations within these specific sites of the RNAPC are likely to be strongly adaptive within the 400 ml LTSP populations. A similar trend is observed at 40 ml. The majority of RNAPC adaptations occurring within the 40 ml LTSP populations occur within one of only four specific sites of the enzyme complex. Clones mutated within one of these sites tend to appear in multiple independently evolving 40 ml LTSP populations. Again, this suggests that these four sites of the RNAPC are adaptive within the 40 ml populations. However, intriguingly the four sites of the RNAPC that are convergently mutated at 400 ml differ from those convergently mutated at 40 ml (**Figure 4**). It, therefore, seems that while the RNAPC is involved in adaptation to LTSP under all volumes, different sites of the enzyme complex may be involved in adaptation at 400 ml compared to 40 ml.

The RNAPC is responsible for all cellular transcription. Therefore, it seems reasonable to propose that adaptations within the RNAPC are likely to affect gene expression. The RNAPC emerges as a target for adaptation in numerous evolutionary experiments, in which *E. coli* was exposed to such diverse selective pressures as heat shock^42^, carbon starvation^43^, antibiotic exposure^44^, heavy metal exposure^45^ and resource exhaustion^7,46,47^. Different sites of the enzyme complex appear to be involved in adaptation to the various selective pressures. We show here that the sites of the RNAPC involved in adaptation can vary even between very similar experiments, in which only a single variable (growth volume) is altered. Why would different sites of the RNAPC tend to be involved in adaptation to different conditions, including conditions that are quite similar? Mutations to different sites of the RNAPC are likely to affect gene expression differently and the precise changes to gene expression which are adaptive are likely to vary between selective pressures. Furthermore, in addition to varying in their adaptive effects, mutations within different sites of the RNAPC likely also vary in their unintended consequences. We have previously demonstrated that the 400 ml LTSP RNAPC adaptations are highly antagonistically pleiotropic, in that they reduce the ability of bacteria to grow within fresh LB^7^. A similar analysis we carried out on the 40 ml RNAPC adaptations, reveals that they also display similar effects of reducing the ability of the cells carrying them to grow within fresh LB (**Figure S1**, Materials and methods). Since the RNAPC is involved in the transcription of all genes, it is quite likely that a mutation to the RNAPC that benefits growth under a particular condition by changing gene expression in a certain way, may also change the expression of other genes in ways that are less desirable. This likely explains these mutations’ antagonistically pleiotropic effects. In order for changes to a specific site of the RNAPC to be adaptive under a specific condition, such undesirable pleiotropic effects need to minimally manifest under that condition. It is thus possible that specific sites are involved in adaptation under specific conditions, because mutations within them change gene expression in the desired manner while minimizing adverse pleiotropic effects on gene expression. The fact that a relatively minor change in conditions leads to adaptation through such different mutations to the RNAPC may suggest that RNAPC adaptations are extremely sensitive to the conditions imposed. It is possible that under more complex conditions, such as those imposed in more natural environments, adaptations through changes to the RNAPC would not be commonly possible.

Combined, our results demonstrate that despite the fact that we observe very convergent dynamics of adaptation under LTSP, it is possible to substantially alter these dynamics by manipulating just a single parameter of the experiment. Certain themes remain consistent across volumes, such as adaptation through the accumulation of genetic alterations, strong positive selection, highly convergent adaptation, and the identity of some of the loci at which adaptation occurs. However, we also observe substantial differences, in the rates of mutation accumulation, in the intensity of positive selection affecting mutation accumulation, and in the adaptations themselves. The effects of altering a single parameter of our evolutionary experiment are likely not unique to this evolutionary experimental setup. By varying parameters of other evolutionary experiments, we should also be able to understand which elements of the experiment can be easily manipulated and which remain stable to minor perturbations.

## Acknowledgments

This work was supported by an ISF grant (No. 756/17, to RH) and by the Rappaport Family Institute for Research in the Medical Sciences (to RH). The described work was carried out in the Rachel & Menachem Mendelovitch Evolutionary Process of Mutation & Natural Selection Research Laboratory.

## References

1. Zambrano, M. M. & Kolter, R. Escherichia coli mutants lacking NADH dehydrogenase I have a competitive disadvantage in stationary phase. J. Bacteriol. 175, 5642–5647 (1993).

2. Zambrano, M. M. et al. Microbial competition: Escherichia coli mutants that take over stationary phase cultures. Science (80-.). 259, 1757–1760 (1993).

3. Finkel, S. E. & Kolter, R. Evolution of microbial diversity during prolonged starvation. Proc. Natl. Acad. Sci. 96, 4023–4027 (1999).

4. Liao, C.-H. & Shollenberger, L. M. Survivability and long-term preservation of bacteria in water and in phosphate-buffered saline*. Lett. Appl. Microbiol. 37, 45–50 (2003).

5. Finkel, S. E. Long-term survival during stationary phase: evolution and the GASP phenotype. Nat. Rev. Microbiol. 4, 113–120 (2006).

6. Rozen, D. E., Philippe, N., Arjan de Visser, J., Lenski, R. E. & Schneider, D. Death and cannibalism in a seasonal environment facilitate bacterial coexistence. Ecol. Lett. 12, 34–44 (2009).

7. Avrani, S., Bolotin, E., Katz, S. & Hershberg, R. Rapid Genetic Adaptation during the First Four Months of Survival under Resource Exhaustion. Mol. Biol. Evol. 34, 1758–1769 (2017).

8. Farrell, M. J. & Finkel, S. E. The Growth Advantage in Stationary-Phase PhenotypeConferred by rpoS Mutations Is Dependent on the pH andNutrientEnvironment. J. Bacteriol. 185, 7044–7052 (2003).

9. Battesti, A., Majdalani, N. & Gottesman, S. The RpoS-Mediated General Stress Response in Escherichia coli. Annual Review of Microbiology 65, 189–213 (2011).

10. Kram, K. E. & Finkel, S. E. Culture Volume and Vessel Affect Long-Term Survival, Mutation Frequency, and Oxidative Stress of Escherichia coli. Appl. Environ. Microbiol. 80, 1732–1738 (2014).

11. Riedel, T. E., Berelson, W. M., Nealson, K. H. & Finkel, S. E. Oxygen consumption rates of bacteria under nutrient-limited conditions. Appl. Environ. Microbiol. 79, 4921–4931 (2013).

12. Kram, K. E. et al. Adaptation of Escherichia coli to Long-Term Serial Passage in Complex Medium: Evidence of Parallel Evolution. mSystems 2, 1–12 (2017).

13. Kram, K. E. & Finkel, S. E. Rich Medium Composition Affects Escherichia coli Survival, Glycation, and Mutation Frequency during Long-Term Batch Culture. Appl. Environ. Microbiol. 81, 4442–4450 (2015).

14. Wein, T. & Dagan, T. The effect of population bottleneck size and selective regime on genetic diversity and evolvability in bacteria. Genome Biol. Evol. 11, 3283–3290 (2019).

15. Baym, M. et al. Inexpensive multiplexed library preparation for megabase-sized genomes. PLoS One 10, 1–15 (2015).

16. Deatherage, D. E. & Barrick, J. E. Identification of Mutations in Laboratory-Evolved Microbes from Next-Generation Sequencing Data Using breseq. in 165–188 (2014). doi: 10.1007/978-1-4939-0554-6_12

17. Maurer, L. M., Yohannes, E., Bondurant, S. S., Radmacher, M. & Slonczewski, J. L. pH Regulates Genes for Flagellar Motility, Catabolism, and Oxidative Stress in Escherichia coli K-12. J. Bacteriol. 187, 304–319 (2005).

18. Slonczewski, J. L., Rosen, B. P., Alger, J. R. & Macnab, R. M. pH homeostasis in Escherichia coli: measurement by 31P nuclear magnetic resonance of methylphosphonate and phosphate. Proc. Natl. Acad. Sci. 78, 6271–6275 (1981).

19. Weiner, L. & Model, P. Role of an Escherichia coli stress-response operon in stationary-phase survival. Proc. Natl. Acad. Sci. 91, 2191–2195 (1994).

20. Booth, I. R. Regulation of cytoplasmic pH in bacteria. Microbiol. Rev. 49, 359–78 (1985).

21. Zilberstein, D., Agmon, V., Schuldiner, S. & Padan, E. Escherichia coli intracellular pH, membrane potential, and cell growth. J. Bacteriol. 158, 246–52 (1984).

22. Deatherage, D. E. & Barrick, J. E. Identification of Mutations in Laboratory-Evolved Microbes from Next-Generation Sequencing Data Using breseq. in 165–188 (2014). doi: 10.1007/978-1-4939-0554-6_12

23. Wen-Hsiung L., G. D. Fundamentals of Molecular Evolution. 2nd Edition. Q. Rev. Biol. 77, 57–57 (2002).

24. Holmqvist, E., Li, L., Bischler, T., Barquist, L. & Vogel, J. Global Maps of ProQ Binding In Vivo Reveal Target Recognition via RNA Structure and Stability Control at mRNA 3′ Ends. Mol. Cell 70, 971-982.e6 (2018).

25. Zhang, M. et al. Structures of the Escherichia coli PutA Proline Dehydrogenase Domain in Complex with Competitive Inhibitors †, ‡. Biochemistry 43, 12539–12548 (2004).

26. Sasaki, H., Sato, D. & Oshima, A. Importance of the High-Expression of Proline Transporter PutP to the Adaptation of Escherichia coli to High Salinity. Biocontrol Sci. 22, 121–124 (2017).

27. Kim, Y.-M. et al. Purification, Reconstitution, and Characterization of Na+/Serine Symporter, SstT, of Escherichia coli. J. Biochem. 132, 71–76 (2002).

28. Campbell, J. W. & Cronan, J. E. Escherichia coli FadR Positively Regulates Transcription of the fabB Fatty Acid Biosynthetic Gene. J. Bacteriol. 183, 5982–5990 (2001).

29. Lioliou, E. E. et al. Phosphorylation activity of the response regulator of the two-component signal transduction system AtoS–AtoC in E. coli. Biochim. Biophys. Acta - Gen. Subj. 1725, 257–268 (2005).

30. Ferrández, A., García, J. L. & Díaz, E. Transcriptional Regulation of the Divergent paa Catabolic Operons for Phenylacetic Acid Degradation in Escherichia coli. J. Biol. Chem. 275, 12214–12222 (2000).

31. Gottesman, S., Clark, W. P. & Maurizi, M. R. The ATP-dependent Clp protease of Escherichia coli. Sequence of clpA and identification of a Clp-specific substrate. J. Biol. Chem. 265, 7886–7893 (1990).

32. Butland, G. et al. Interaction network containing conserved and essential protein complexes in Escherichia coli. Nature 433, 531–537 (2005).

33. Sniegowski, P. D., Gerrish, P. J. & Lenski, R. E. Evolution of high mutation rates in experimental populations of E. coli. Nature 387, 703–705 (1997).

34. Giraud, A. Costs and Benefits of High Mutation Rates: Adaptive Evolution of Bacteria in the Mouse Gut. Science (80-.). 291, 2606–2608 (2001).

35. LeClerc, J. E., Li, B., Payne, W. L. & Cebula, T. A. High Mutation Frequencies Among Escherichia coli and Salmonella Pathogens. Science (80-.). 274, 1208–1211 (1996).

36. Gross, M. D. & Siegel, E. C. Incidence of mutator strains in Escherichia coli and coliforms in nature. Mutat. Res. Lett. 91, 107–110 (1981).

37. Voordeckers, K. et al. Adaptation to High Ethanol Reveals Complex Evolutionary Pathways. PLOS Genet. 11, e1005635 (2015).

38. Mehta, H. H. et al. The Essential Role of Hypermutation in Rapid Adaptation to Antibiotic Stress. Antimicrob. Agents Chemother. 63, 1–18 (2019).

39. Raynes, Y., Wylie, C. S., Sniegowski, P. D. & Weinreich, D. M. Sign of selection on mutation rate modifiers depends on population size. Proc. Natl. Acad. Sci. 115, 3422–3427 (2018).

40. Wielgoss, S. et al. Mutation rate dynamics in a bacterial population reflect tension between adaptation and genetic load. Proc. Natl. Acad. Sci. 110, 222–227 (2013).

41. Good, B. H. & Desai, M. M. Evolution of Mutation Rates in Rapidly Adapting Asexual Populations. Genetics 204, 1249–1266 (2016).

42. Tenaillon, O. et al. The molecular diversity of adaptive convergence. Science (80-.). 335, 457–461 (2012).

43. LaCroix, R. A. et al. Use of adaptive laboratory evolution to discover key mutations enabling rapid growth of Escherichia coli K-12 MG1655 on glucose minimal medium. Appl. Environ. Microbiol. 81, 17–30 (2015).

44. Degen, D. et al. Transcription inhibition by the depsipeptide antibiotic salinamide A. Elife 2014, 1–29 (2014).

45. Graves, J. L. et al. Rapid evolution of silver nanoparticle resistance in Escherichia coli. Front. Genet. 5, 1–13 (2015).

46. Nandy, P., Chib, S. & Seshasayee, A. A Mutant RNA Polymerase Activates the General Stress Response, Enabling Escherichia coli Adaptation to Late Prolonged Stationary Phase. mSphere 5, 1–16 (2020).

47. Chib, S., Ali, F. & Seshasayee, A. S. N. Genomewide Mutational Diversity in Escherichia coli Population Evolving in Prolonged Stationary Phase. mSphere 2, 1–15 (2017).

