## Supplementary material for "Culture volume influences the dynamics of adaptation under long-term stationary phase": Figure S1

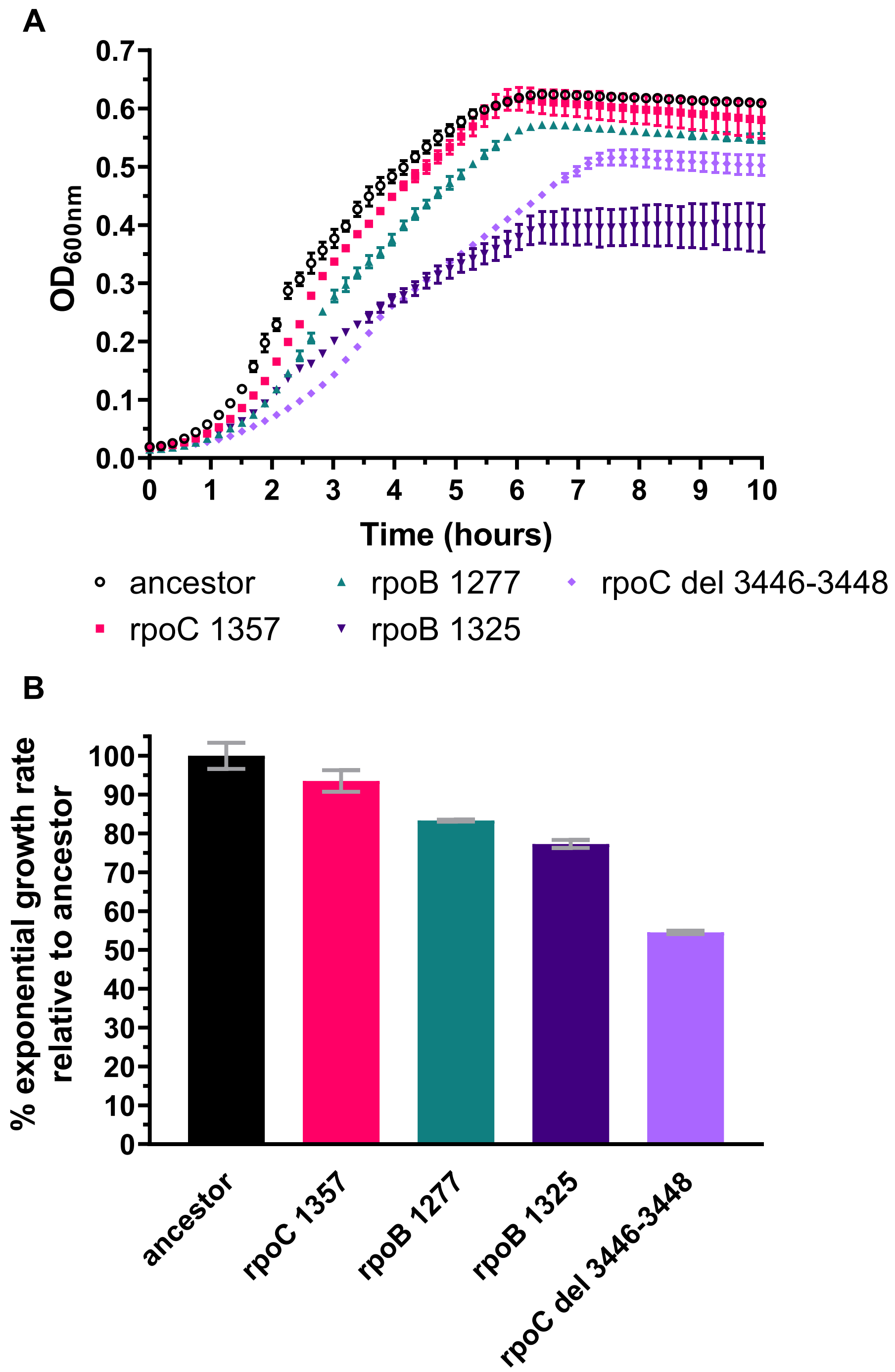


**Supplementary Figure S1. The 40 ml RNAPC convergent adaptations are associated with a cost to growth within fresh LB** (A) Growth curves - Clones carrying each of the convergent 40 ml RNAPC adaptations that were otherwise identical to the ancestral wildtype strain were grown in a 96 well plate within a plate reader and OD600 was measured every 10 minutes. For comparison sake the ancestral strain was also similarly monitored. Two independent clones from each strain were analyzed and for each clones two independent experiments were carried out. For each strain, depicted is the mean OD 600 values, across the four independent experiments, during the first ten hours of growth as a function of time. Error bars represent standard deviations around these means. (B) Mean exponential growth rates of each 40 ml clone tested, relative to the mean exponential growth rate of the ancestral (wildtype) *E. coli* strain (Materials and methods).
